# Single-cell RNA sequencing of murine liver reveals an aligned circadian clock and cell-population specific circadian regulated pathways

**DOI:** 10.1101/2025.03.24.644906

**Authors:** Anthony Veltri, Manabu Nukaya, Ksenija Korac, Patrick B. Schwartz, Sean M. Ronnekleiv-Kelly

## Abstract

The circadian clock is tightly connected to metabolism, which is evident in various metabolic processes performed by the liver. Perturbation of these processes due to circadian dysregulation leads to liver specific pathology. The liver is composed of multiple different cell populations each with distinct functions contributing to organ homeostasis, but individual cell population contributions to circadian clock function is not yet known. Single-cell RNA sequencing provides the opportunity to understand clock function and oscillating gene expression within an organ system at the individual cell population level that would allow for better understanding of the crosstalk between the circadian clock and metabolic pathways within the liver. In the past, barriers to achieving this goal included complexity associated with generating single-cell RNA sequencing time series data as well as the complexity of data analysis. Here, we established a protocol that enabled the generation of murine liver cell population time series data, as well as a methodological approach to evaluate the core molecular clock and oscillating gene expression in individual cell populations. Using a combination of normalized coefficient of variation, clock-correlation and aggregate pseudobulk, we found a robust and aligned circadian clock in each of the cell populations. We then employed a pseudoreplicate / pseudobulk strategy to identify oscillating gene expression and benchmarked against bulk RNA sequencing data; we demonstrated that many metabolic genes were oscillating in several of the cell populations, including non-hepatocyte clusters. Finally, we identified oscillating genes unique to specific cell populations that play critical roles in liver function. The findings in this study lay an important foundation for understanding clock function and contributions of oscillating gene function at the individual cell population level in liver.

## Introduction

The circadian clock is a unique biological system that maintains homeostasis by coordinating the timing of essential processes across the 24-hr day/night cycle. This organization is achieved by a series of interlocked transcriptional-translational feedback loops (TTFL) comprised of circadian activators (positive arm) and circadian repressors (negative arm) that generate an autoregulatory cycling of core clock genes (1). In the positive arm, transcription factors CLOCK and BMAL1 form a complex that binds to specific cis-acting E-box elements, promoting the transcription of target repressor genes such as *PER1-3*, *CRY1-2*, and *NR1D1-2*. The protein products of the repressor arm inhibit the activity of the CLOCK-BMAL1 complex either directly via post-translational mechanisms (e.g., CRY, PER) or through suppression of *BMAL1* expression (e.g., NR1D1) (1). This interplay yields rhythmic expression of the molecular clock over 24 hours, and maintains cellular and organ homeostasis in the setting of environmental cycles (e.g., feeding-fasting cycle) and systemic factors (2–4).

The fundamental role of the clock is in metabolic processes, which is evident from numerous studies in fatty acid metabolism, *de novo* lipogenesis, glycogen metabolism, and insulin / insulin-like growth factor signaling, among others (4–8). This intrinsic link between the circadian clock and metabolic processes consequently underlies the strong connection between circadian dysregulation and metabolic pathologies, many of which are a result of liver dysfunction (6, 9). These include obesity, diabetes, metabolic dysfunction associated liver disease (MALD) and liver cancer, which reflects the principal metabolic role of the liver (6–11). Investigation of liver specific clock function has provided much insight into circadian clock regulated pathways, and how perturbation of these pathways leads to liver-specific pathology. For instance, circadian disruption caused by mutation in core clock genes (e.g., *Per1^-/-^;Per2^-/^*) or through chronic circadian misalignment (jetlag) results in profound rewiring of liver metabolic processes including lipid deposition, hepatic triglyceride and free fatty acid accumulation, altered glucose and glycogen metabolism, imbalanced bile acid synthesis, and dysregulation of *Car* (*Nr1i3*) that ultimately yields hepatocellular carcinoma (9).

Landmark studies such as these have used elegant whole organ analyses combined with hepatocyte-specific mutant models (e.g., *Alb^Cre^;Bmal1^fx/fx^*) to determine the crosstalk between the (dysregulated) circadian clock and metabolic pathways within the liver (5, 8, 9). However, the liver is composed of several different cell populations, each serving distinct functions contributing to organ homeostasis (12–14). For instance, cell populations outside of hepatocytes contribute significantly to liver function and pathologic states, such as Kupffer cells and endothelial cells (12, 13, 15). Additionally, hepatocytes can be further sub-divided into discrete populations (e.g., pericentral hepatocytes, periportal hepatocytes, etc.) and each sub-population may manifest differences in clock regulated pathways (e.g., zonation of bile acid biosynthesis in hepatocytes) (14). Yet, the role of the molecular clock in specific cell populations within the liver has been challenging to assert. Various strategies employed are cell autonomous core clock gene manipulation (e.g., hepatocyte knock-out models), use of immortalized cells or isolation of specific cell populations *ex vivo* (e.g., antibody-bead sorting) (5, 8). However, these strategies may prove challenging for exploring sub-population specific clock function and diurnal (oscillating) gene expression in a complex tissue such as liver.

Single-cell RNA sequencing provides the opportunity of understanding clock function within an organ system at the resolution of individual cell populations. A barrier to achieving this goal is the complexity associated with generating single-cell RNA sequencing time series data from mouse liver. In fact, at present, there exist few mammalian single-cell RNA sequencing time-course studies that investigate clock function including one in mouse dermal cells and a separate in mouse SCN (8, 16–18). Further, there is the added complexity of data analysis since many techniques applicable for bulk RNA sequencing are less effective when analyzing single-cell data (17). Consequently, we present here the first single-cell RNA sequencing time course study (to our knowledge) in mouse liver with samples obtained every four hours. We test the hypothesis that the circadian clock is aligned in each of the different cell populations, and that oscillating gene expression varies depending on the cellular context (including differences among the hepatocyte population). We found that the circadian clock was robust and aligned in each of the cell populations. We assessed key metabolic pathways that occur in liver and found that many of the metabolic genes were oscillating in several of the cell populations, including non-hepatocyte clusters (e.g., endothelial cells). Further, we identified cell population specific oscillating genes that play critical roles in liver function. Given the importance of the circadian clock and diurnal physiology in liver health and liver pathologic states (e.g., metabolic dysfunction associated steatohepatitis) (3, 5, 7–9, 19), this investigation is meant to serve as a critical foundation for future work.

## Results

### The core molecular clock is intact in each cell population

To evaluate circadian clock function in individual cell populations, we established a protocol that enabled generation of liver single cell suspensions at four-hour intervals. We performed retrograde *in vivo* liver perfusion by cannulating the vena cava, followed by removal of perfused liver, dissociation into single cell suspension, measurement of cell viability (to ensure greater than 90% viability) (20), and cell fixation; this was done every four hours for twenty-four hours. The fixed cell suspensions containing parenchymal and non-parenchymal cells were then sequenced and annotated for cell clustering and analysis. Cell types identified were concordant with prior hepatic single-cell studies, including multiple sub-populations of hepatocytes and distinct subclusters of endothelial cells (**Fig 1A**, **C and S1 Fig**) (8, 12, 14). Roughly the same number of cells were acquired at each Zeitgber Time (ZT), with similar cell distribution by ZT (**Fig 1B**, **D and S2 Fig**). In total, there were 34,921 cells over six timepoints.

**Fig 1.**
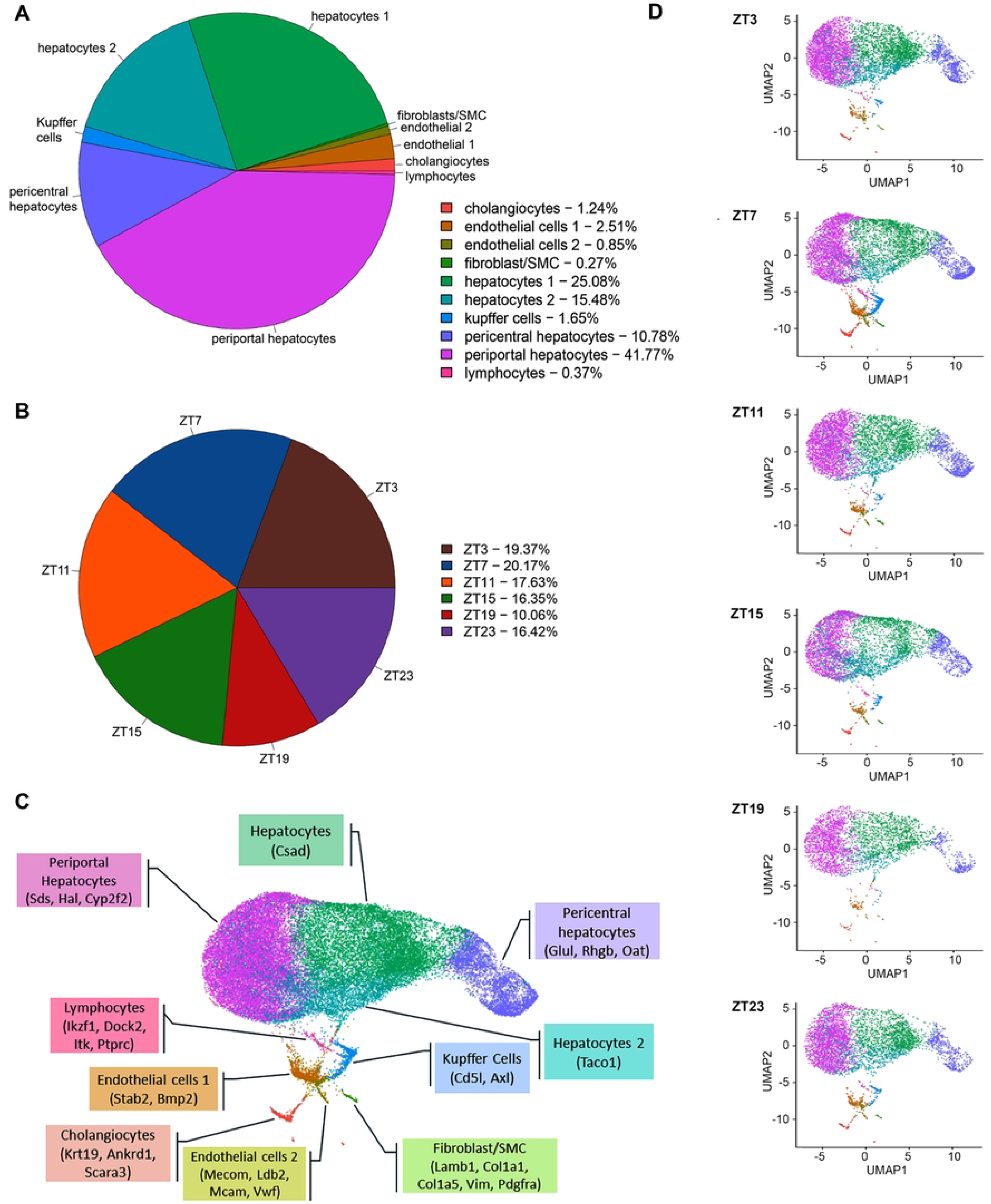
Single-cell RNA sequencing of the whole murine liver cells across 24hr. Eight-week-old C7BL/6J mice were entrained to standard lighting conditions (6am/6pm) with ad libitum access to food and water. Starting at 9am (ZT3), one mouse was euthanized every four hours over a 24-hour period [1pm (ZT7), 5pm (ZT11), 9pm (ZT15), 1am (ZT19) and 5am (ZT23)] and livers collected for single-cell RNA sequencing. Both parenchymal cells (PCs) and non-parenchymal cells (NPCs) were isolated from each harvested liver by a two-step perfusion-dissociation method and displayed ∼90% viability prior to the cell fixation. A total of 1 million PCs and NPCs cells were fixed using the Evercode™ Cell Fixation v3 kit (Parse Biosciences, Seattle WA). Single cell libraries were constructed and eight sublibraries were pooled and sequenced on an Illumina NovaSeq X Plus sequencer using a 10B flow cell. An average sequencing depth was ∼180k reads per cell. Depicted in **A)** is a pie chart illustrating the average cell population composition and quantity isolated from all liver samples (n=6) across the 24-hour period. The 10 identified cell populations represent both PCs and NCs with majority of the populations being PCs. **B)** Pie chart illustrating the percentage of cells in each individual time point across the 24-hour period. The number of cells across all time points was comparable. **C)** Uniform manifold approximation and projection (UMAP) plot visualization of the 10 distinct liver cell clusters that include all cells from each of the time points combined into a single projection. Each cell cluster is labeled with the established gene markers that were used to annotate the clusters listed in the parenthesis. **D)** UMAP plot visualization of the 10 distinct liver cell clusters by individual time points. All cell clusters are well represented across each time point without preferential isolation in one specific time point compared to the others.

We assessed circadian clock function in each cell population by normalized coefficient of variation (nCV), clock correlation, and gene expression across 24 hours. These complementary approaches enable a measure of circadian clock health and robustness (21–24). We first employed nCV to gain an understanding of the relative amplitude of core clock gene expression in the single cell dataset. We compared the nCV of core clock genes from the entire dataset (i.e., inclusive of all cell populations) to our prior work evaluating bulk RNA sequencing in a similarly metabolic organ, the mouse pancreas (**S3 Fig**) (25). In the analysis of pancreatic cells, each of the core clock genes were shown to be rhythmic on meta2d analysis (25). Presently, we found a strong concordance between ‘whole’ liver and pancreas (no significant difference in mean nCV), indicating robust oscillation of core clock genes in the liver. We then evaluated nCV for each individual cell population, which demonstrated variability depending on the cell population (**Fig 2 and S4 Fig**). For instance, nCV ranged from 0.359 – 1.536 for the core clock genes in fibroblasts (even though oscillations looked greater, this is *normalized* within the cell population) versus 1.126 – 6.015 in periportal hepatocytes. However, this was not indicative of a ‘weaker’ clock and instead likely represents cell-population specific variations or alternatively the differences in cell number per population (i.e., hepatocyte populations comprise greater cell numbers). Evidence supporting this assertion is the fact that the phase correlation of the core clock genes were highly preserved (clock correlation matrix (24)) and core clock genes demonstrated rhythmic expression for each cell population (**Fig 2 and S5 Fig**). Therefore, while the nCV analysis may have been skewed by cell number, the combined analysis of core clock gene function was robust to differences in cell numbers (e.g., fibroblasts vs periportal hepatocytes) and revealed an intact circadian clock in each cell population.

**Fig 2.**
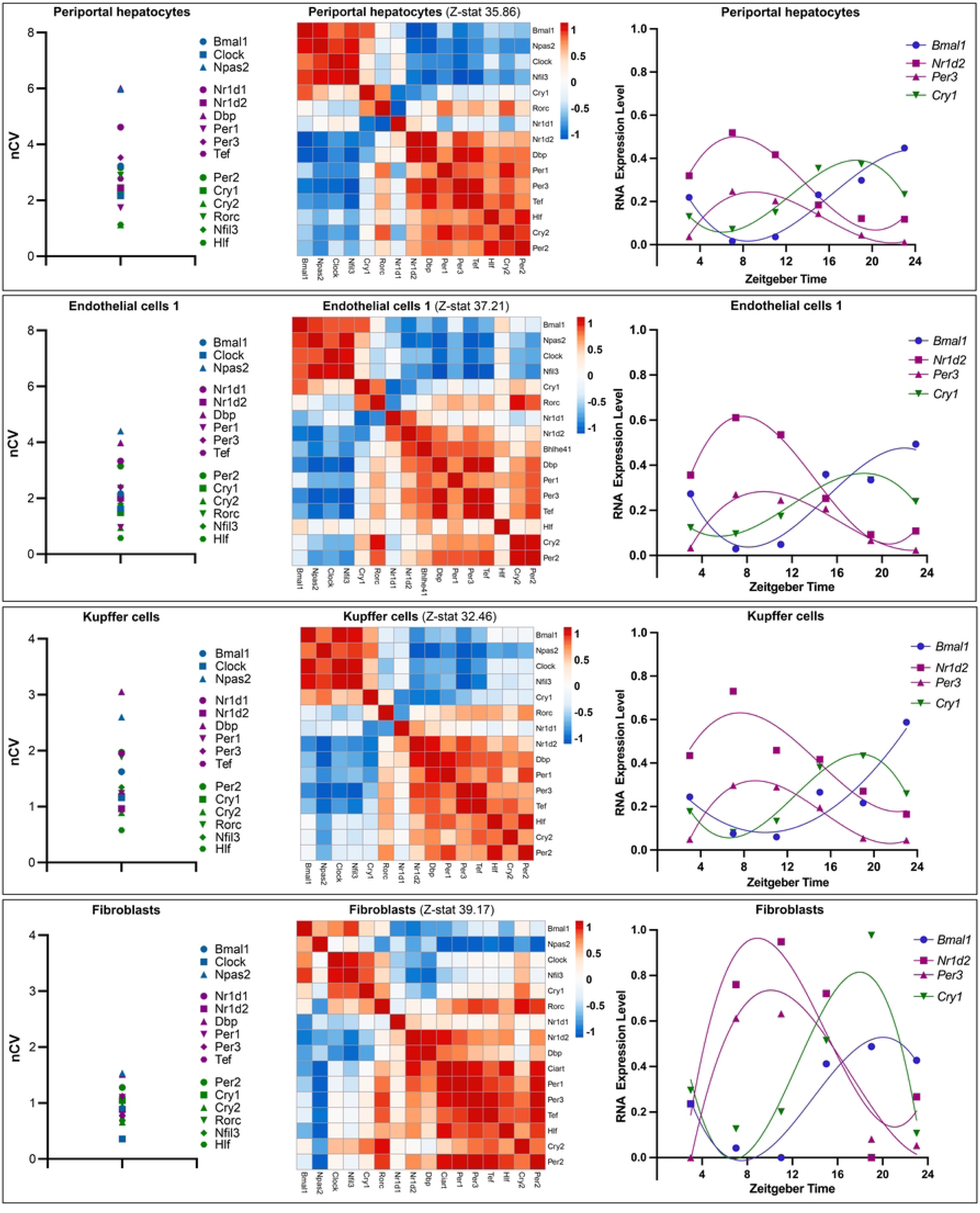
Cell population specific evaluation of circadian clock function. Shown is the normalized coefficient of variation (nCV), clock correlation matrix, and gene expression level over time in four separate cell populations including periportal hepatocytes, endothelial cells, Kupffer cells and fibroblasts. Four separate cell populations were evaluated for clock function including periportal hepatocytes, endothelial, Kupffer cells and fibroblasts. (**Left**) The normalized coefficient of variation (nCV) tightly correlates with the normalized relative amplitude (rAMP) of core clock gene expression. rAMP is a critical feature of rhythmically expressed genes. Evaluated clock genes were categorized as E-Box activators (blue), early E-Box targets (purple), and E-Box and other inputs (green). All four representative cell populations demonstrate a robust clock. (**Middle**) Shown is the clock correlation which corresponds to the phase relationship of core clock genes. For instance, the positive arm of the clock (e.g., BMAL1, CLOCK) drives transcription of the negative arm of the clock (e.g., PER, CRY). Thus, when *BMAL1* expression peaks, *CLOCK* expression should also peak (Spearman’s rho (ρ) ∼1), and negative arm members (*e.g*., *PER1-2*) should be at a trough (ρ ∼ -1). Using the set of core clock genes, the correlation of each gene against the others is determined and mapped in matrix form. Z-statistic (z-stat) values represent the extent that these established correlations are preserved in each of the cell populations evaluated, with higher values indicating phase relationships that are closer to the circadian gene atlas baseline (i.e., highly preserved clock correlation). (**Right**) Core clock gene expression levels are depicted for each cell population over 24 hours, pertaining to the liver samples collected at different Zeitgeber Times (ZT). The RNA expression level is aggregate RNA expression from the cell population at each ZT. Shown is the oscillation of the selected core clock genes (e.g., *Bmal1, Nr1d2, Per3* and *Cry1*), which comprise genes from each of the categories (e.g, E-box activators, etc.). This was curve fit using a third-order polynomial curve fit and allows for visualization of the expected phase relationships between core clock genes (e.g., *Bmal1* and *Nr1d2* are anti-phasic).

The pseudobulk approach was used to assess core clock gene expression in each cell population (**Fig 2**). This approach aggregates cells to yield a single biological replicate for each cell population at each ZT, and ensures high biological accuracy from single-cell data (17, 26). Genes from each of the categories including E-box activators (*Bmal1*, *Clock*, *Npas 2*), strong E-box target genes (*Nr1d1*, *Nr1d2*, *Dbp*, *Per1*, *Per3*, *Tef*), and those with E-box targets plus other inputs (*Per2*, *Cry1*, *Cry2*, *Rorc*, *Nfil3*, *Hlf*) demonstrated clear rhythmic expression in each of the cell populations evaluated. The expected phase relationships were also conserved within each cell population. For instance, anti-phasic expression of positive arm members (e.g., *Bmal1*) and negative arm members (e.g., *Nr1d2, Per3*) was evident. Additionally, the phase relationships appeared conserved across the cell populations with synchronous peak phase and trough of expression. To investigate further, we examined core clock gene expression overlayed on the UMAP embedding, and found that this pattern held true for each of the core clock genes including E-box activators, strong E-box targets and those with E-box plus other inputs (**Fig 3** and **S6 Fig**) (27). Peak phase was then mapped onto a phase diagram with three separate diagrams pertaining to the core clock gene categories (e.g., E-box activators); this revealed that the core clock genes showed aligned expression across cell populations (i.e., narrow window of peak phase of expression). Finally, the expression pattern (i.e., peak and trough) of the core clock genes across 24 hours was also strongly aligned in each population of cells (**Fig 4**). This confirmed synchrony of the core molecular clock in cell populations acquired from *in vivo* liver tissue.

**Fig 3.**
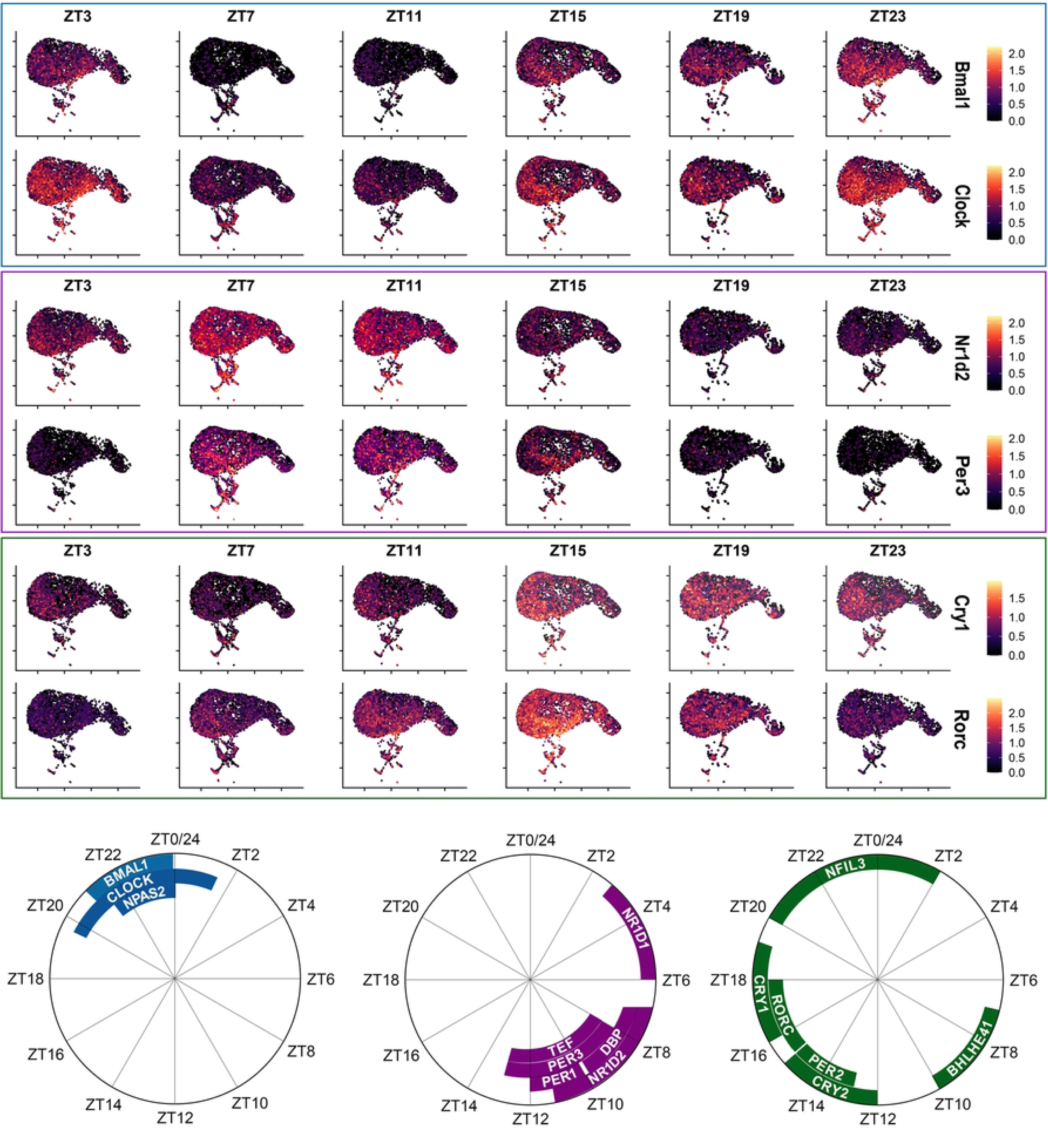
Peak phase of expression of core clock genes. Shown in the top panel is a Pseudotime trajectory visualizing the expression of the representative individual genes from the E-Box activators (*Bmal1*, *Clock*), strong E-Box targets (*Nr1d1*, *Per3*), and genes with E-Box plus other inputs (*Cry*, *Rorc*) across the different time points. Cells are plotted in order of expression (i.e., higher expression cells are plotted over cells of lower expression). Additional core clock genes are shown in Supplemental Figure 4. Expression levels at each time point encompasses all identified liver cell populations with lowest expression demonstrated by dark coloration and highest expression represented by lighter coloration. Zero pseudotime is set to the point of the lowest gene expression. The aforementioned genes display variation in expression between different time points demonstrating rhythmicity that is concordant in all identified cell populations. At the bottom is a graphical representation of the peak phase of expression for each circadian category: E-Box activators (blue), strong E-Box early targets (purple), and genes with E-Box plus other inputs (green) in all samples (n=6) across the 24-hour period. The phase distribution of each bar (gene) represents the peak phase in expression among the various cell populations; thus, a narrower span of ZT indicates a tighter range of peak expression.

**Fig 4.**
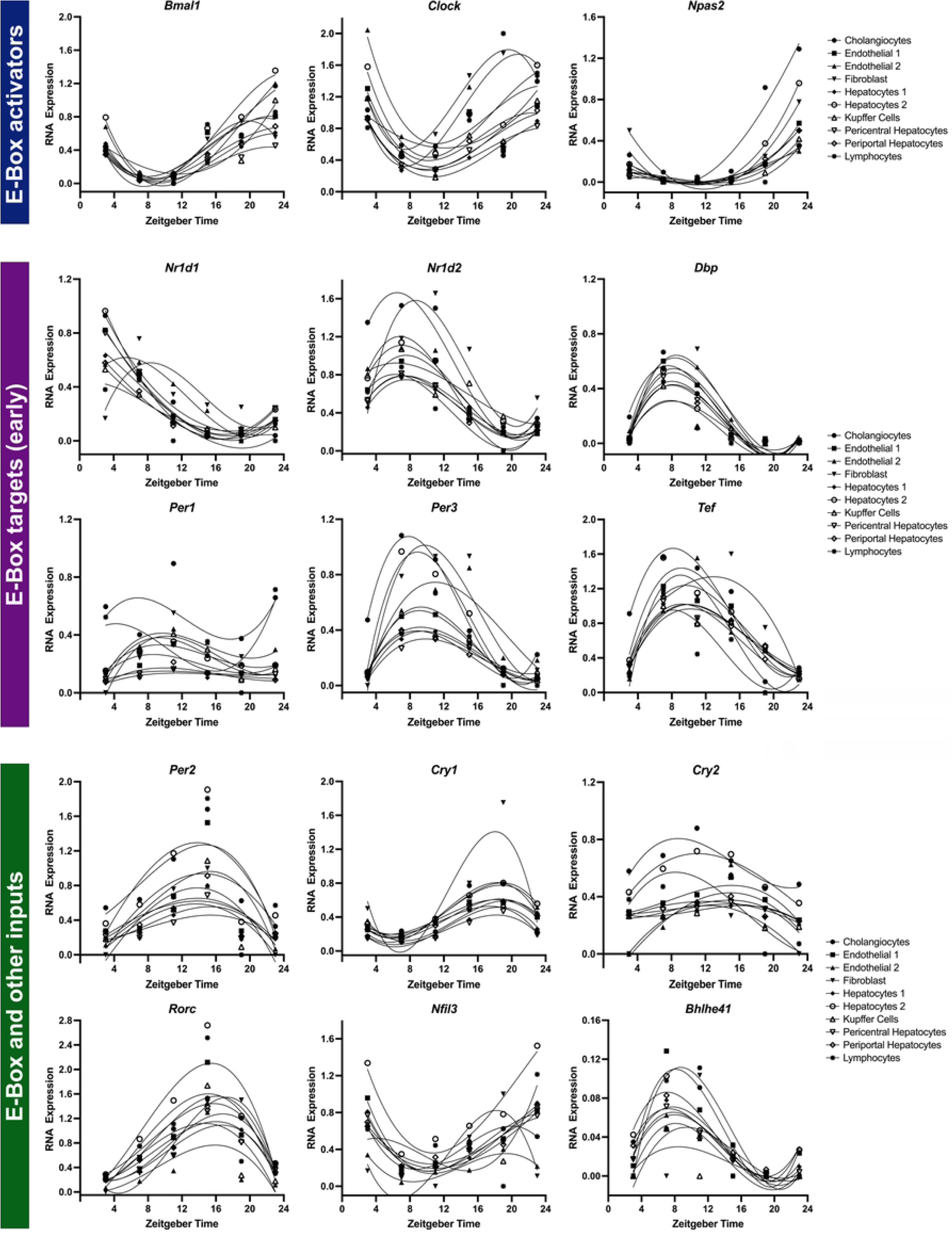
Phase distribution plots of the core clock and clock associated genes. Core clock and clock associated gene expression levels are depicted at different Zeitgeber Times (ZT) for each liver cell population over 24 hours. At the top (blue) are the E-Box activators (*Bmal1*, *Clock*, *Npas2*). BMAL1-CLOCK proteins and BMAL1-NPAS2 proteins bind to the E-Box elements and activate the expression of the other core clock and clock associated genes. Those genes in the middle are primarily driven by E-Box activation (E-Box early targets, purple) which include *Nr1d1*, *Nr1d2*, *Dbp*, *Per1*, *Per3*, and *Tef*. Their peak phase of expression comes up almost immediately after *Bmal1* as expected. At the bottom are genes with E-Box and other inputs (green) which include *Per2*, *Cry1*, *Cry2*, *Rorc*, *Nfil3*, and *Bhlhe41*, and whose peak phase of expression is typically observed at the later Zeitgeber time points. This data demonstrates preserved phase distribution in each of the cell populations, with evidence of robustly oscillating gene expression (curve fit using a third-order polynomial curve fit). Additionally, the different cell populations are closely correlated in terms of the peak phase of expression of different core clock and clock associated genes.

### Single-cell RNA sequencing strategy for detecting diurnal gene expression

After establishing an intact molecular clock, we next evaluated for diurnal (oscillating) gene expression (outside of core circadian clock genes) in each of the liver cell populations. This is a challenging endeavor given the high variability in expression patterns amongst single-cells and considering that most methods have been built for bulk RNA sequencing data (17, 28). For instance, certain methods cannot accommodate uneven replicates across time points, which is pertinent for single-cell RNA sequencing data given the variable number of cells obtained for each cell population at each timepoint. Further, treating each cell as a replicate may increase the false positive rate and incorrectly identify oscillating genes (17, 26). To minimize false positives, pseudobulk data is preferential for asserting oscillating gene expression, but may be limited by false negatives (16–18). We applied ARSER, JTK-cycle, Lomb-Scargle and MetaCycle meta2d analysis to the pseudobulk data and identified only few rhythmically expressed genes in any cluster using a false-discovery rate *q* < 0.05 (**S1 File**) (29). Furthermore, despite strong evidence of an intact circadian clock in each of the cell clusters (**Fig 2-4**), none of the core clock genes demonstrated rhythmicity (*q* < 0.05) via pseudobulk approach.

Our assertion was that the pseudobulk approach was too stringent in identifying oscillating genes with only one ‘biological replicate’ per timepoint. To probe further, we examined known clock driven genes that had previously been identified in mouse liver (30). This group of clock driven genes was established through elegant studies in wild-type, *Bmal1* mutant and *Cry1/2* mutant mice, under ad-libitum and night-restricted feeding paradigms (bulk RNA sequencing) (30). Out of 529 clock driven genes identified from this bulk RNA sequencing work, we selected the top 100 genes and ordered them by phase of expression based on the meta2d phase. We then used this ordered list and evaluated gene expression over time (depicted by heat map) for each cell cluster (**Fig 5A and S7 Fig**). This demonstrates an apparent oscillating pattern of expression with discrete peak and trough phases, which would be expected for clock-driven genes. To further strengthen the support for rhythmic gene expression, we plotted peak phase of expression of the clock driven genes between the bulk RNAseq dataset and our single-cell dataset and found a strong correlation (Pearson r = 0.8619, *p* < 0.0001) (**Fig 5B**). When limiting the correlation analysis to core clock genes, the phase correlation (Pearson r = 0.9700, *p* < 0.0001) (**Fig 5C**) and overlap in peak phase (**S8 Fig**) were even more robust. Taken together, this supported that the few identified rhythmically expressed genes when applying the pseudobulk method to single-cell data was likely limited by one biological replicate per time point; the trade-off of higher biological accuracy is detection of fewer cycling genes with pseudobulking (17).

**Fig 5.**
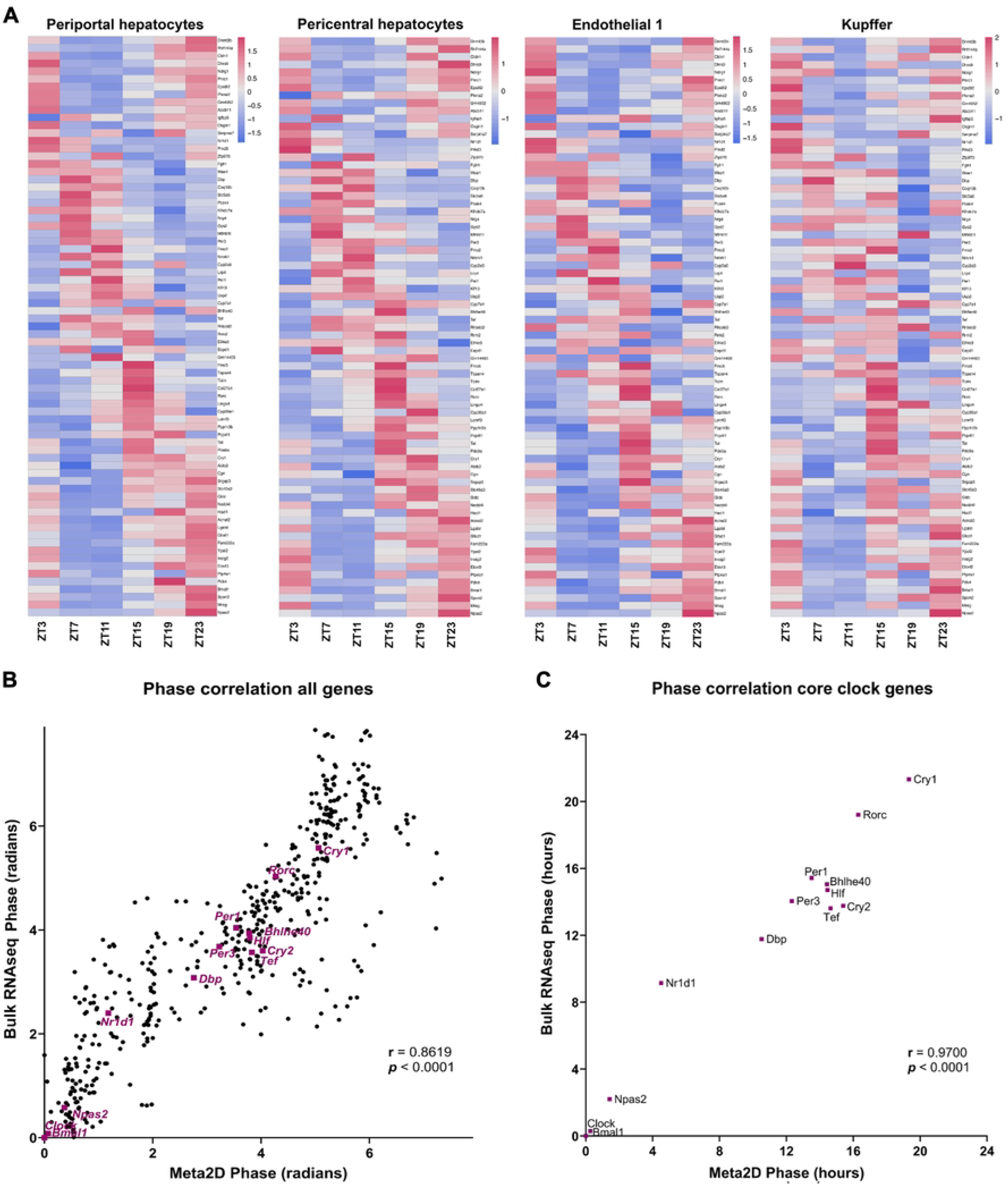
Phase correlation of clock driven genes. **A)** The top 100 clock-driven genes were identified from an existing bulk liver RNA sequencing dataset (30). These genes were ordered in the single-cell dataset according to phase of expression (meta2d) using average pseudobulk and gene expression depicted for each ZT. This was visualized via heatmap for representative cell populations. **B)** The 529 clock driven genes from murine liver^24^ were plotted to evaluate the Pearson r correlation between the bulk RNA phase and single-cell phase (Meta2d phase). This revealed a strong phase correlation for all 529 genes (r = 0.8619, p < 0.0001) and **C)** an even stronger correlation for the core clock genes (r = 0.9700, p < 0.0001).

To overcome this limitation, we reexamined the data by treating each cell as a biological replicate. Although each cell is not truly independent within a cell cluster (26), this approach is commonly utilized in single-cell analysis (17). Using this strategy, we identified a substantial increase in number of oscillating genes in each cluster via meta2d analysis (FDR *q* < 0.05) (**S2 File**). For instance, in the pericentral hepatocytes population, this included the core clock genes as well as many other putative oscillating genes (e.g., 3490 genes identified as oscillating). This does raise concern for a high false positive rate, which is a known limitation of this approach (17). Moreover, while we were able to complete this analysis for many of the cell clusters including those with low numbers of cells (underscoring an advantage), this approach was not feasible for the larger cell clusters including periportal hepatocytes (14585 cells across all time points) and hepatocytes 1 (8759 cells across all time points), as well as the entire population of cells. Feasibility was limited by computation time, which precluded completion of convergence in these larger cell clusters (analysis did not converge by 15 days), which was also reported by other investigators (17). This highlights another significant limitation of treating each cell as a biological replicate, particularly as single-cell data sets increase in cell number and depth of sequencing.

To achieve a balance between feasibility, biological accuracy and detection of oscillating genes, and to address the present limitations, we employed a pseudoreplicate / pseudobulk strategy (**S9 Fig**) (17, 31). We randomly partitioned each population of cells (each cluster) into three subsets and applied the pseudobulk method to each subset. This resulted in three pseudoreplicates which can be treated as biological replicates per cell population at each ZT; MetaCycle meta2d was then applied to identify rhythmically expressed genes. To improve rigor of the approach and minimize false positive detection, the pseudoreplicate / pseudobulk strategy was conducted for ten separate trials in each cell population (**S3 File**) (17). Each trial of random partitioning and pseudobulking was followed by detection of rhythmically expressed genes (meta2d *q* < 0.05). Gene rhythmicity detected in at least half of the iterations should be sufficient for considering oscillation (particularly with *q* < 0.05); for instance, in endothelial cells 1 cluster, *Bmal1, Cry1, Nr1d1-2, Per2-3, Rorc,* and *Tef* were rhythmic in 10/10 iterations, but *Dbp* (9), *Nfil3* (7) and *Cry2* (1) were rhythmic in fewer than ten iterations. The advantage for a lower iteration threshold (e.g., five out of ten iterations versus ten out of ten iterations) is detection of an increased number of rhythmically expressed genes (**S10 Fig and S4 File**). However, outside of the core clock genes, the most confident oscillating genes would demonstrate rhythmicity in all ten iterations. Therefore, to minimize false positives, we included those genes with a threshold of *q* < 0.05 and ten out of ten iterations, which still resulted in a significant number of oscillating genes and clock-driven genes identified in most clusters (**S11 Fig**).

### Metabolic genes oscillate in several cell clusters

The cell cluster specific identification of oscillating genes revealed that many of the rhythmically expressed genes were found in more than one population of cells (**S11 Fig**). Further, many of the oscillating genes identified in the cell clusters were known clock-driven genes (**Fig 6A and S12 Fig**), such as elongation of very long-chain fatty acids 3 (*Elovl3*), which is involved in hepatic lipogenesis, or *Insig2*, the protein product of which regulates SREBP1 and consequently genes involved in *de novo* lipogenesis, such as *Elovl5, Elovl6, Fasn* and *Acly* (**Fig 6B and Fig 7A**) (30, 32). Interestingly, these circadian regulated genes that are critical for hepatic *de novo* lipogenesis were consistently oscillating in the endothelial population, and not just the hepatocytes. This was also observed for genes involved in glycogen regulation, such as glycogen synthase 2 (*Gys2*), which was rhythmically expressed in endothelial cells, kupffer cells and hepatocyte populations (**Fig 7B**). Meanwhile, oscillation of *Slc2a2* (*Glut2*) was restricted to the hepatocyte populations, as were genes involved in NAD+ metabolism (*Nampt* and *Nmrk1*), supporting that circadian regulation of hepatic glucose export and NAD+ salvage are mediated by the hepatocyte populations (**Fig 7B-C**). Further, the peak phase of expression of these genes involved in hepatic metabolic processes was aligned in each of the cell populations and was concordant with previously published work (7, 11). To investigate the pathways associated with oscillating genes within each population, we performed pathway analysis in the various cell clusters with sufficient rhythmically expressed genes, including endothelial cells 1, hepatocytes 1, hepatocytes 2, pericentral hepatocytes and periportal hepatocytes (**S5 File**). Examining the top ten pathways by KEGG and Reactome analysis revealed several overlapping pathways that were primarily metabolic (e.g., PPAR signaling, Fatty acid metabolism) and related to bile acid metabolism (**S13-15 Figs**).

**Fig 6.**
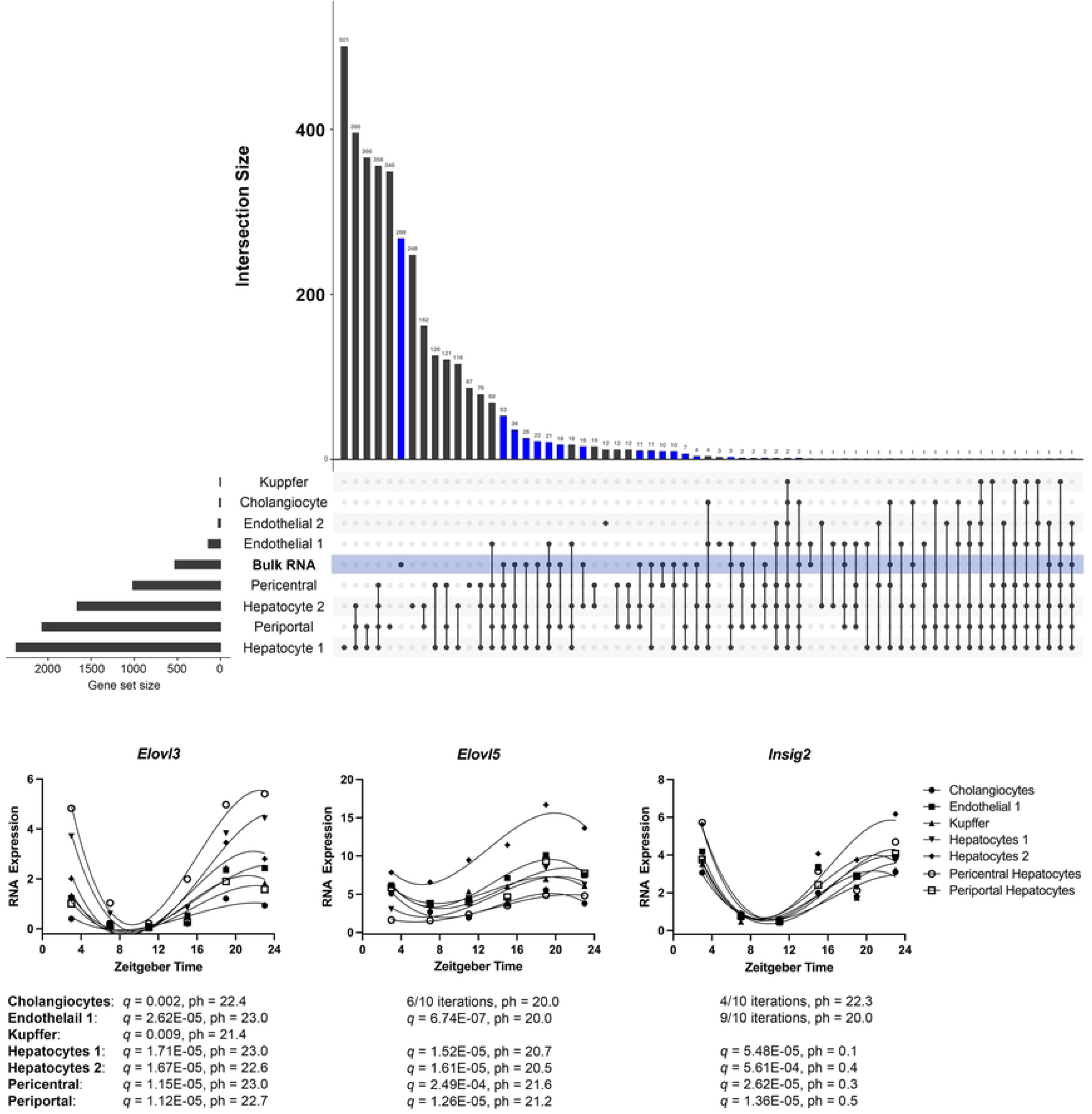
Oscillating genes by cell cluster. **A)** Oscillating genes in each individual cell population were determined by performing pseudoreplicates combined with pseudobulking, followed by meta2d analysis (rhythmically expressed: FDR *q* < 0.05). Shown is an UpSet Plot with the number of oscillating genes in each cell population (gene set size) and the number of intersecting genes between the cell populations. Intersecting genes pertain to the number of oscillating genes that are expressed by two or more cell populations (dots and line), or the number of genes that are solely expressed by the individual cell cluster (dot). Depicted in blue are the known clock-driven genes determined by bulk RNA sequencing analysis of liver from wildtype and *Bmal1*-mutant or *Cry1*/*Cry2*-mutant mice (bulk RNA) (30). **B)** Shown are three genes involved in hepatic lipogenesis that were found to be oscillating (*q* < 0.05) in multiple cell types. For each cell type, the FDR *q*-value is depicted (or the number of iterations in which *q* was < 0.05) along with the peak phase of expression (ph). *Elovl3* and *Insig2* were also identified as clock-driven genes from the bulk RNA sequencing investigation.

**Fig 7.**
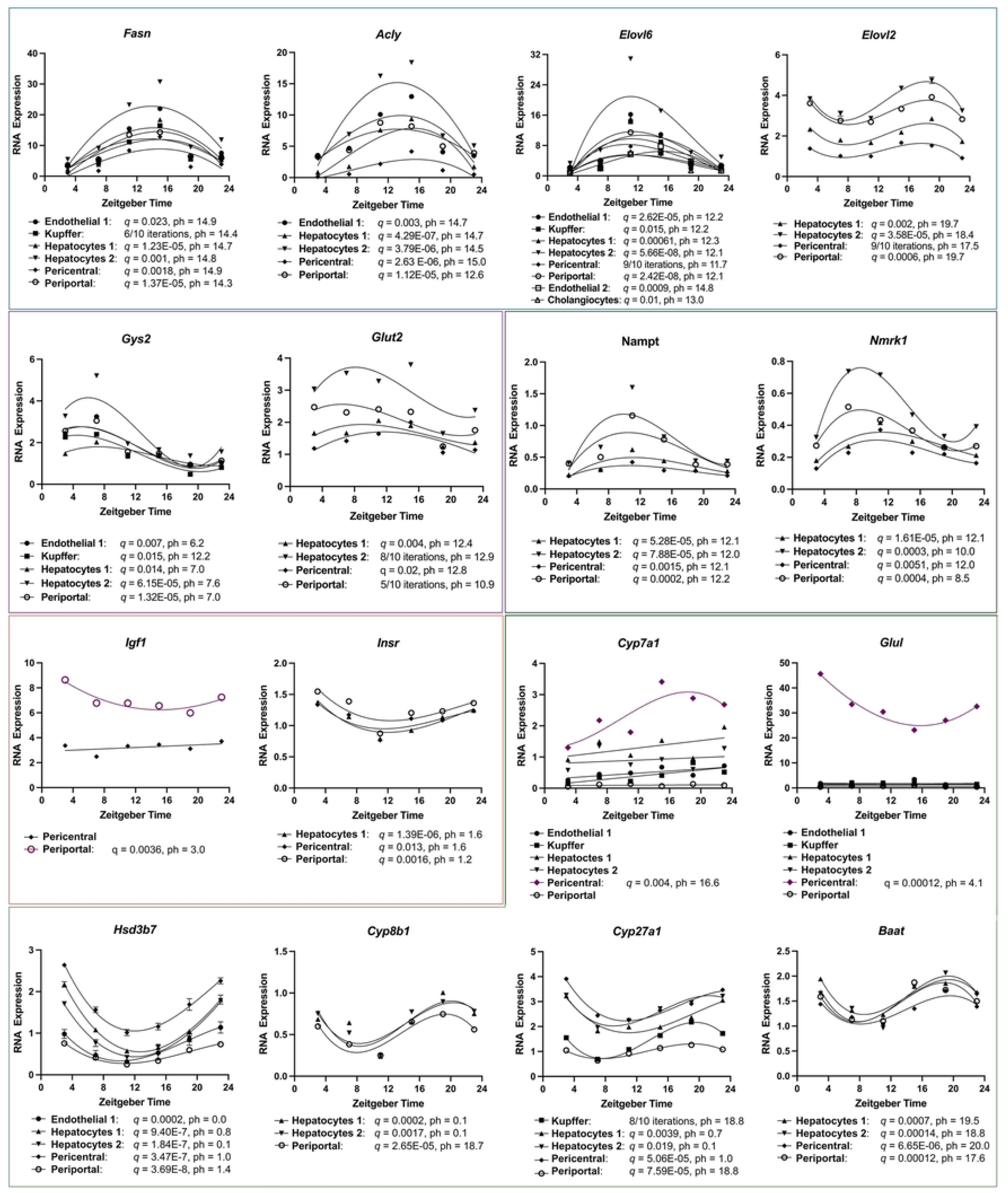
Metabolic genes in liver cell populations. Genes involved in metabolic pathways that were found to be oscillating in various cell populations are depicted. Listed below the graph is the individual cell population with FDR *q*-value (or number of iterations that it was found to be oscillating) and peak phase of expression. Shown are genes involved in (**A**) *de novo* lipogenesis, (**B**) glycogen and glucose regulation, (**C**) NAD+ salvage pathway, (**D**) insulin-like growth factor signaling, and (**E**) bile acid synthesis. In the bile acid synthesis box, glutamine synthase (*Glul*) is also depicted; this gene helps to define the pericentral hepatocyte population (annotation) and was also found to be oscillating.

### Pathway analysis highlights cell cluster specific oscillating gene expression

There were also many distinct rhythmically expressed genes in each cell population that were not detected in other cell clusters, indicating that each cell cluster has unique sets of rhythmic gene expression (**Fig 6A**). For instance, insulin and insulin-like growth factor 1 (*Igf1*) are powerful systemic modulators of the circadian clock, strongly inducing *Per* transcription and PER translation, thereby synchronizing cellular clocks with feeding (4). While the insulin receptor, *Insr* (shown in prior work to be rhythmically expressed in liver) (5, 8, 33), was rhythmically expressed in periportal hepatocytes, hepatocytes 1 and pericentral hepatocytes, *Igf1* was only oscillating in the periportal hepatocyte population (**Fig 7D**). Previously, *Igf1* was shown to be highly expressed in the periportal layers, which aligns with our finding that *Igf1* is oscillating only in the periportal cells (14). This circulating growth factor and feeding regulated hormone therefore appears to interface with the circadian clock by oscillatory expression in the periportal hepatocyte subset (4, 14), which is notable given that periportal hepatocytes ‘face’ the portal venous inflow where dietary nutrients are transported (14). In turn, IGF-1 can bind its receptors such as *Insr* (IR), which was found to be rhythmically expressed in phase with *Igf1*, to transmit the signal from feeding to entrainment of the circadian clock (4).

Also zonally distributed is the bile acid biosynthesis pathway, where the first step is regulated by *Cyp7a1*, converting cholesterol to 7-alpha-hydroxycholesterol in pericentral hepatocytes (14, 34). The pericentral hepatocytes are distinguished by *Glul* (glutamine synthetase) expression, which interestingly we also found was oscillating. Bile acid synthesis follows a progression from pericentral hepatocytes to periportal hepatocytes (location of bile acid conjugation and secretion) through an enzymatic cascade (*Hsd3b7*→*Cyp8b1*→*Cyp27a1*→Baat) (34). *Cyp7a1* is a known clock-driven gene (30) and remarkably was found to only be rhythmically expressed in the pericentral hepatocytes; the other enzymes involved in bile acid synthesis were oscillating in all four hepatocyte populations. The importance of this observation is that *Cyp7a1* is the rate limiting step for bile acid biosynthesis, uncovering a hepatocyte sub-population specific link between the circadian clock and regulation of a major metabolic pathway in the liver (**Fig 7E**).

This shows that different cell populations within the liver express discrete oscillating genes throughout a 24-hour period that contribute to diverse essential functions. In support, our pathway analysis (**Fig 8**) demonstrated unique pathways for individual cell populations, such as ‘KEAP1-NFE2L2 pathway’ in pericentral hepatocytes, ‘degradation of cysteine and homocysteine’ in the periportal hepatocytes population, ‘Toll Like Receptor TLR1:TLR2 cascade’ in hepatocytes 2 population, and ‘regulation of insulin-like growth factor (Igf) transport and uptake by insulin-like growth factor binding proteins (Igfbps)’ in Endothelial 1. Moreover, the periportal hepatocyte population exhibited higher expression of pathways involved in respiratory electron transport, ATP production and oxidative phosphorylation (**S15 Fig**). These pathways containing oscillating gene expression highlight the functions of these cell populations, such as the pericentral hepatocytes that are farthest removed from oxygenated blood (i.e., higher oxidative stress relative to periportal hepatocytes), or that IGF is a circulating growth factor within the vascular space (requiring uptake and transport by endothelial cells), or the well-oxygenated location of periportal hepatocytes allowing for ATP-demanding tasks (4, 14). Determining the comprehensive set of genes and pathways that are under circadian control and altered by circadian dysregulation will be examined in future work, but the present study is an important foundation for understanding clock function and contributions of oscillating gene function at the individual cell population level in liver.

**Fig 8.**
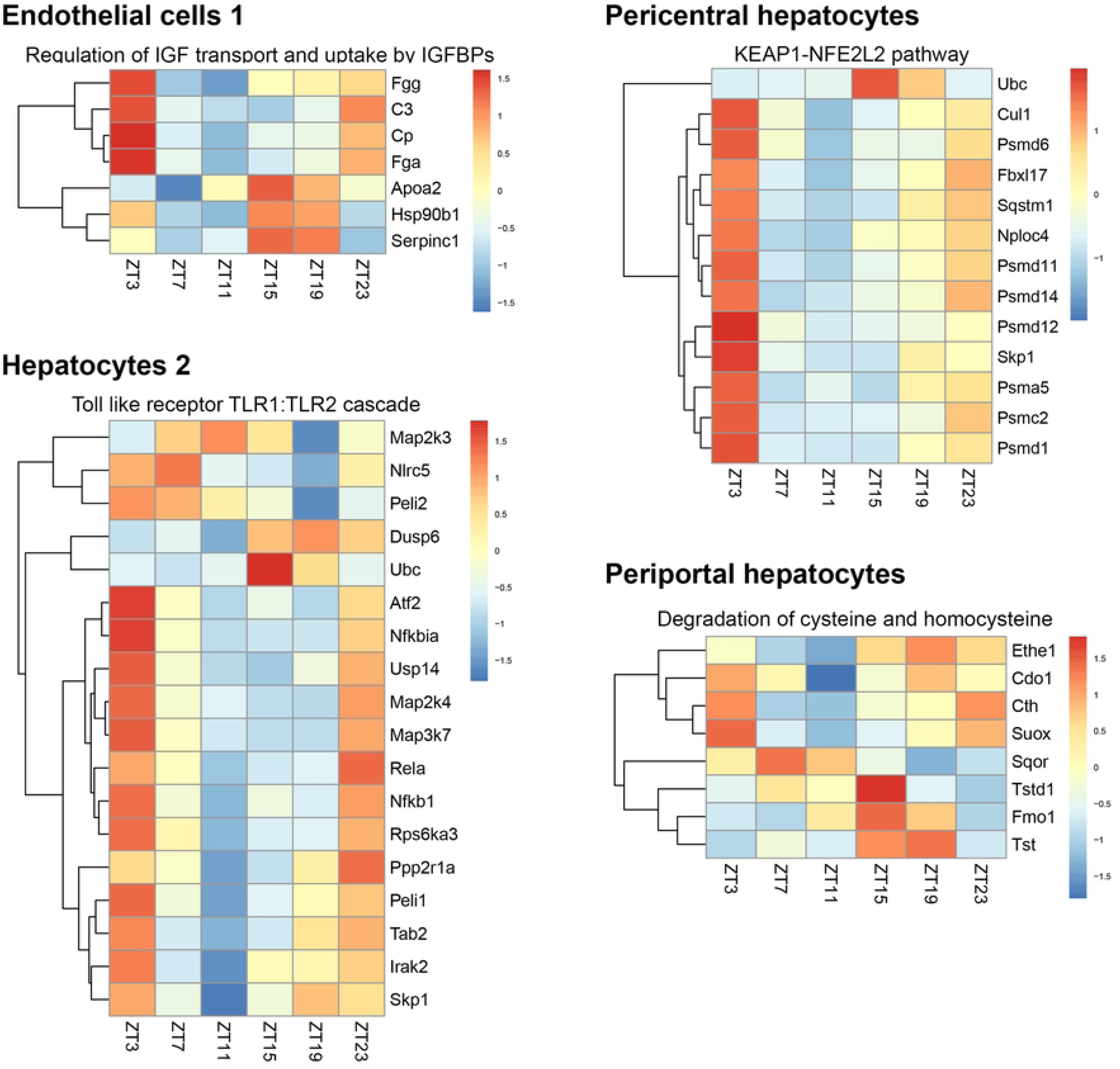
Pathway analysis of oscillating genes in discrete cell clusters. Rhythmically expressed genes were determined for each cell cluster via a pseudoreplicate / pseudobulk approach. These identified genes were evaluated using Reactome and KEGG pathway analysis. While many of the pathways were overlapping for the various cell clusters (as expected), there were also pathways that were unique to individual cell populations. Depicted are the heatmaps representing some of these pathways, showing the gene expression over time of the oscillating genes that comprised these pathways. For instance, the KEAP1-NFE2L2 pathway in pericentral hepatocytes contains several genes involved in ubiquitin dependent proteolysis, and nearly all oscillate with a peak phase of expression at ZT3, indicating an important timing of this pathway in this cell cluster.

## Discussion

In the current study, we performed liver single-cell RNA sequencing at four-hour intervals over a 24-hour period. Previous work has examined individual cell populations using single-cell sequencing either as an isolated time point (14), or through isolation of specific populations followed by bulk RNA sequencing (8). We examined the molecular clock function and oscillating gene expression in individual cell populations from murine liver, including four separate hepatocyte populations. This was revealing considering the unique oscillating genes, circadian-driven genes and pathways associated with hepatocyte sub-populations. There was also a substantial overlap in various metabolic processes (e.g., fatty acid metabolism). We assessed roughly 6000 cells for each time point to achieve deep sequencing for each cell (∼197,000 reads / cell). Given the significant variability in expression patterns amongst cells within a cellular subtype (stochastic expression), our approach balanced the identification of individual cell populations with the a priori goal of identifying oscillating gene expression (17, 35).

We used methods established in bulk RNA sequencing (e.g., nCV and clock correlation) and single-cell analysis (pseudobulking) to determine that the circadian clock was intact and aligned in each individual cell population. This would be presumed but has not been shown previously in this number of cell populations in liver (to our knowledge) (8, 16, 18). To benchmark our work, we examined the phase distribution identified in our individual cell populations compared to prior published work of core clock gene expression (bulk RNA) in liver tissue (8, 10, 27). For instance, investigators evaluating clock function in individual liver cell populations isolated hepatocytes (parenchymal cell fraction) and two non-parenchymal cell populations consisting of endothelial cells and Kupffer cells; these were isolated from non-parenchymal cell fraction using anti-CD146 microbeads and anti-F4/80 microbeads, respectively. After isolation every six hours, cells were lysed and bulk RNA sequencing performed to evaluate core clock gene expression, which revealed robust oscillation of core clock genes in hepatocytes and endothelial cells, with lower amplitude rhythmic expression in Kupffer cells (8). Liver demonstrated peak expression of *Bmal1* at ZT22-ZT2, *Cry1* at ZT18-22, *Nr1d1* roughly ZT4-8, *Nr1d2* roughly ZT12-16, and *Per1* around ZT10 (8, 10). This matched up precisely with our single-cell population specific phase distribution of core clock gene expression, and demonstrates that ‘clock timing’ of one population can be extrapolated to other populations under standard conditions in mice. However, the effect of various environmental perturbations (e.g., misalignment or mis-timed feeding) on cell population specific clock phases is yet to be determined, and may impact cell population subsets differently.

Beyond the core clock genes, the presence of significant noise within single-cell RNA sequencing data (i.e., dropouts or high cell-to-cell variability even within individual cell populations) and the paucity of robust methods for single-cell data analysis (rhythmicity) led to significant challenges with identifying oscillating gene expression (17, 28, 35). We ultimately used the approach of pseudobulking to minimize the false positive detection of oscillating genes that may occur (e.g., artifactual elevated expression at a single timepoint), alongside pseudoreplicate generation to mitigate the consequence of false negatives (17). We again benchmarked our data against work by other investigators that had identified clock-driven genes in the liver of mice; this was accomplished by bulk liver RNA sequencing in wildtype mice, *Bmal1*-mutant mice and *Cry1*/*Cry2*-mutant mice. And while we did identify significant overlap (particularly with hepatocyte clusters) with the clock-driven genes, there were also numerous other oscillating genes identified in each cell cluster (e.g., 894 genes in the pericentral hepatocytes cluster that were not amongst the clock-driven genes).

Rhythmicity of gene expression can arise from clock-driven genes, clock-associated genes or rhythmicity induced by systemic cues (e.g., food intake) (10, 30). Thus, only a fraction of rhythmically expressed genes in organs are regulated by the core molecular clock. This is consistent with previous work that has elegantly depicted the myriad circadian clock genes, clock-associated genes and clock-regulated genes in mouse liver (30). Outside of the 529 clock-driven genes identified, there were 190 oscillating genes found to be partially modulated by the clock and 903 oscillating genes driven by systemic cues (1622 total oscillating genes), demonstrating that many of the oscillating genes within the liver are regulated by fasting-feeding cycles or other systemic cues, which has been shown in several studies (8, 10, 30). Concordantly, the majority of rhythmically expressed genes in the liver lose rhythmicity with arrhythmic feeding (10). This is due to the powerful cue of rhythmic food intake, where roughly 80% of food intake occurs during the active phase of mice fed ad-libitum (10). This contrasts with the few hundred genes identified that were specifically regulated by the cell autonomous clock in liver. Thus, although we were able to define core clock gene expression in each of the cell populations, our examination of rhythmically expressed genes does not translate to clock-controlled genes in each of these cell populations. Deciphering clock regulated genes / pathways versus those regulated by systemic cues will be further explored with time-restricted feeding paradigms and core-clock knockout models combined with our single-cell approach.

Another notable aspect of this work is the demonstrated ability to identify oscillating gene expression in specific population subsets within the liver. This extends beyond prior work investigating whole-organ contributions through bulk RNA sequencing. For instance, elongation of very long-chain fatty acids 3 (*Elovl3*) is a known clock-regulated gene and important for hepatic lipogenesis, lipid droplet formation in liver, and diet-induced obesity (32, 36). Notably, this gene was found to be oscillating in all hepatocyte populations, cholangiocytes, endothelial cells and Kupffer cells. Other members of the *Elovl* family, such as *Elovl5* and *Elovl6*, were also found to be oscillating in cholangiocytes, endothelial cells, Kupffer cells (*Elovl6* only) and all hepatocyte populations (7, 19). *Elovl5* is strongly induced by obesogenic diet, mediating *de novo* lipogenesis within the liver and is regulated by *Srebp1* / *Insig2* (7). Concordant with *Elovl5* expression, *Insig2* was found to be rhythmic in cholangiocytes, endothelial cells and all hepatocyte populations. Combined, this indicates that these cell populations may be working in concert with the hepatocytes for *de novo* lipogenesis and lipid droplet formation in the liver. This also supports data demonstrating the involvement of these varied cell populations (Kupffer cells and endothelial cells) in liver metabolic disorders (metabolic dysfunction associated steatohepatitis) (12). Interestingly, another member, *Elovl2*, was only oscillating in the four hepatocyte populations. *Elovl2* has been previously identified as rhythmically expressed in retinal cells and is a strong marker of molecular aging, where methylation of *Elovl2* promoter (causing suppressed expression) increases with age, leading to age-related macular degeneration (and is exacerbated by diabetes) (37, 38). *Elovl2* encodes for a protein that regulates levels of polyunsaturated fatty acids that protect against oxidative stress, suggesting a putative connection between the circadian clock and stress-induced molecular aging specifically within the hepatocyte populations. This could have implications for maintaining a ‘healthy liver clock’ to combat against age-linked or stress-induced diseases such as metabolic syndrome and cancer (39).

Bile acid metabolism is another key pathway in the liver. Bile serves several critical functions in the body such as facilitating absorption of fat and fat soluble vitamins, or as a route for cholesterol excretion (cholesterol is initial substrate for bile acids) (34). Notably, dysregulation of bile acid synthesis can lead to many different pathologies such as gallstones, liver inflammation, atherosclerosis or neurologic disorders (34). Further, bile acids activate various nuclear receptors in the liver that are critical for normal function, such as farnesoid X receptor (FXR). Generation of bile acids from cholesterol is catalyzed by CYP7A1, which is responsible for the first and rate limiting step in bile acid synthesis (generation of 7α-hydroxycholesterol from cholesterol), comprising the ‘gatekeeper’ for 75% of total bile acids produced (40). *Cyp7a1* is most abundant in the pericentral hepatocytes (14), has been identified as a clock-driven gene (30), and circadian misalignment results in altered expression (9). In fact, a key finding in recent work showed that circadian misalignment resulted in suppression of FXR with altered expression of *Cyp7a1*, coupled with dysregulation of bile acid synthesis, cholestasis and development of metabolic dysfunction associated liver disease (9). The dysregulation of bile acids additionally activated *Car* (*Nr1i3*), which promoted development of hepatocellular carcinoma. Remarkably, we found that *Cyp7a1* is only oscillating in the pericentral hepatocytes, strongly suggesting a cell-population specific connection to clock regulation of bile acid production, and to circadian dysregulation-induced liver pathology. We also found that *Nr1i3*, which has previously been reported as an oscillating gene in the liver of mice (and thought to be a clock-controlled gene) (9), is oscillating in specific hepatocyte subsets including periportal hepatocytes, hepatocytes 2, and pericentral hepatocytes.

The level of resolution presented here (e.g., *Cyp7a1*) exemplifies how use of our approach enables dissection of these individual cell populations with respect to diurnal physiology, and in future work, circadian regulated pathways. This is consequential because of the significant mechanistic insight achievable into cell population specific circadian regulated and systemic cue regulated pathways in liver, which will be important for combatting liver pathologic states attributable to circadian disruption or mis-timing of cues (9). In turn, this work opens the door for important lines of inquiry including whether there are cell population specific sensitivities to circadian disruption or environmental perturbations of the clock.

For instance, miscues that occur due to circadian misalignment or obesogenic diet may impact sub-populations of cells more than others (e.g., pericentral hepatocytes), which could inform screening strategies, lifestyle interventions or precision therapeutic strategies. At present, this remains uncertain and will be a focus of future work. There are limitations to consider for the current study. For instance, clusters with few cells such as the fibroblasts did not demonstrate rhythmically expressed genes using the pseudoreplicate / pseudobulk approach. We did identify oscillating genes when treating each cell as a biological replicate (e.g., *Insig2*, *Acly*, *Elovl3*, *Elovl6*; **S2 File**), but none of the core clock genes were detected as rhythmic in fibroblasts despite using this approach. This is an inherent limitation of single-cell RNA sequencing data given the stochastic noise associated with single-cell expression (28, 35), and should be an important consideration for time series single-cell data. A strategy to address this limitation would include sequencing of parenchymal cell fractions (i.e., hepatocytes) and non-parenchymal cell fractions separately, to minimize ‘dilution’ of non-parenchymal cell fractions. These fractions are separated during the liver cell dissociation process and thus this strategy can be seamlessly employed in future work. An additional limitation is that only male mice were used in this investigation. Given the precise nature of the experimental protocol to achieve high quality single-cell suspensions every four hours, only one mouse liver was able to be harvested at each time point. Prior work that we benchmarked our data against was performed in male mice (8, 30), which was an important contributing factor for our approach. Future work will need to include female mice given the known sex-dependent differences in oscillating gene expression in individual cell populations from liver, and sex-dependent differences in liver pathology (36, 41). Taken together, this present investigation represents a consequential foundation for future work.

## Materials and Methods

### Parenchymal and non-parenchymal liver cell isolation and fixation

All animal studies were conducted according to an approved protocol (M005959) by the University of Wisconsin School of Medicine and Public Health Institutional Animal Care and Use Committee. Mice were housed in an Assessment and Accreditation of Laboratory Animal Care–accredited selective pathogen free facility (UW Medical Sciences Center) on corncob bedding with chow diet (Mouse diet 9F 5020; PMI Nutrition International) and water ad libitum. Eight-week-old male C57BL/6J mice were housed in standard conditions under a 12-h:12-h light-dark (LD) cycle (Light-on 6 am-6 pm, Light-off 6 pm-6 am). To collect liver tissues, mice were sacrificed at 4-hour intervals over 24 hours corresponding to Zeitgeber time [10 am (ZT4), 2 pm (ZT8), 6 pm (ZT12), 10 pm (ZT16), 2 am (ZT20), 6 am (ZT24)].To isolate parenchymal cells (PCs) and non-parenchymal cells (NPCs) for single-RNA sequencing from mouse liver, we used the two-step liver perfusion-dissociation method described by Wang (20). Mice were anesthetized by isoflurane and cannulated the Internal Vena Cava (IVC) with 26-gauge IV catheter. The liver was perfused in situ through the IVC with EGTA working buffer using a peristaltic pump (3-3.5 mL/min, 5 min) and digested by Pronase working buffer (0.543 mg/mL pronase, perfused for 7 min) and Collagenase D working buffer (0.5 mg/mL collagenase D, perfused for 7 min), respectively. After the digestion, the liver was gently removed and dissociated by shaking with Liver dissociation buffer (0.6 mg/mL pronase, 0.5 mg/mL collagenase D) for 20 min at 39°C. The liver homogenate was filtrated through a 70 mm cell strainer, centrifuged at 580 g for 10 min at 4°C, and then washed with GBSS buffer twice. PCs and NPCs were isolated by density gradient centrifugation using Nycodenz buffer (centrifuged 1,380 g, 17 min, 4°C). After lysis of red blood cells by 1x RBC lysis buffer, PCs and NPCs were collected by centrifuge at 400 g for 5 min at 4°C. The PCs/NPCs number was counted, and cell viability was measured using trypan blue staining (>80% cell viability of PCs is reliable).

### Single-cell RNA sequencing cell fixation

A total of 1 million PCs and NPCs cells (0.3-0.4 million PCs, 0.6-0.7 million NPCs) were immediately fixed using the Evercode Cell Fixation v3 kit (Parse Biosciences, Seattle WA) for single-RNA sequencing. In brief, cells were transferred to a BSA-coated 1.5mL tubes and centrifuged (400xg at 4°C) for 5 minutes. Cells were resuspended in 187.5μL of Prefixation Master Mix. The solution was then filtered into a new tube by using a 70μm cell strainer. 62.5 μL of cell Fixative Master Mix was added to the tube and the solution was mixed by gentle pipetting (3x). The solution was incubated on ice for 10 minutes followed by the addition of 20μL of the Permeabilization solution and incubated on ice again for 3 minutes. 250μL of Fix and Perm Stop buffer was added, mixed gently by pipetting (3x) and centrifuged (400xg at 4°C) for 5 minutes. Cells were resuspended in 100μL of Cell Storage Master Mix and filtered into a new BSA-coated 1.5uL tube by using a 70μm cell strainer. We performed a second Cell Storage Master Mix rinse to ensure better cell retention. Fixed cells were stained by Trypan blue and counted by using a hemocytometer. Fixed cells were stored at -80°C until sequencing.

### Construction of single-cell RNA sequencing libraries

Single-cell libraries were constructed using the Parse Evercode™ WT v3 kit (Parse, Seattle, WA). Six fixed single-cell suspensions were submitted on dry ice to the University of Wisconsin Biotechnology Center, where cell concentrations were measured using the Luna-FX7 Automated Cell Counter (Logos Biosystems, Annandale, VA) with AO/PI staining (Logos Biosystems, Annandale, VA). Following the instructions outlined in the Parse Evercode WT v3 User Manual v1.3, a target of 24,900 cells per sample was loaded onto the Barcoding Round 1 Plate across forty-eight wells, undergoing three rounds of barcoding. After barcoding, cells were divided into eight sublibraries, and sample-specific indexes (Parse UDI Plate-WT) were added via sample index PCR (10 cycles). Following sample index PCR, double-size selection was performed using SPRIselect (Beckman Coulter, Brea, CA), the final libraries were quantified and assessed for size and adapter dimer contamination using the Qubit High Sensitivity DNA Kit (Thermo Fisher Scientific) and Tapestation Systems (Agilent, Santa Clara, CA), respectively. The eight sublibraries were pooled and sequenced on an Illumina NovaSeq X Plus sequencer using a 10B flow cell.

### Single-cell RNA sequencing data analysis

#### Raw data processing

FASTQ files were processed using the Parse Biosciences pipeline version 1.3 to identify cell barcodes and align and count reads. Samples were aggregated using the ‘comb’ pipeline mode.

#### Downstream scRNAseq processing

Gene expression data was then processed using the Seurat version 5 pipeline for data integration (42). Data was loaded into Seurat. Cells were evaluated for quality and thresholds were set for number of UMIs detected (<5000 UMIs) and mitochondrial content (>35% mitochondrial gene content). Mitochondrial content thresholds were set permissively in concordance with previous work that observed high mitochondrial read counts in high-quality hepatocyte cells (43). Cells from outlier barcoding wells were also removed. Doublets were removed with the R package scDblFinder (44). Samples were normalized with SCTransform v2 with 3000 unique features (45). Samples were integrated using Seurat’s RPCA integration (42). A principal component analysis (PCA) was performed with the number of dimensions (principal components) selected based on an elbow plot. Data were visualized using the Uniform Manifold Approximation and Projection for Dimension Reduction (UMAP) (46). Clusters were identified by the Louvain algorithm performed on all cells in the dataset, then cell types were identified by marker expression and applied to the quality-controlled dataset. Each cluster was well represented across all timepoints in the dataset, and gene expression was either summed and divided by total counts, then converted to a CPM-like value using a 1E6 scaling factor or averaged for cells within each timepoint and cluster to perform pseudobulk analysis. For pseudobulk / pseudoreplicate analysis, cells in each cluster were partitioned randomly into three pseudoreplicates and gene expression was averaged within pseudoreplicates. This procedure was repeated for 10 trials.

#### Cycling detection

Meta2D analysis from the MetaCycle package was performed on each cluster (29). Normalized coefficient of variation (nCV) and clock correlation analysis were performed with the nCV package (22). Upset plots for overlapping cycling genes were created using the package UpSetR and venn diagrams were created using the package eulerr (47, 48). Gene enrichment pathway analysis was performed using the package clusterProfiler to compare genes that were identified as cycling in all iterations with gene sets from GO (49–51), KEGG (52), and Reactome (53).

## Acknowledgements

We would like to acknowledge the University of Wisconsin Biotechnology Center Gene Expression Center for their assistance in optimizing the single-cell RNA sequencing fixation protocol.

## Supporting Information

**S1 Fig**. **Single-cell RNA sequencing cell population annotation**. Depicted is the Uniform manifold approximation and projection (UMAP) plot visualizing the expression of the representative individual genes that were used for cell cluster annotation, including identification of pericentral hepatocytes (*Glul*, *Oat*, *Rhbg*), periportal hepatocytes (*Cyp2f2*, *Hal*, *Sds*), hepatocytes 1 (*Csad*), hepatocytes 2 (*Taco1*), general endothelial cells (*Cdh5*, *Pecam1*), endothelial cells 1 (*Bmp2*, *Stab2*), endothelial cells 2 (*Mecom*, *Ldb2*, *Mcam*, *Vwf*), cholangiocytes (*Ankrd1*, *Krt19*, *Scara3*), Kupffer cells (*Cd5l*, *Axl*, *Cd68*, *Itgam*), lymphocytes (*Ikzf1*, *Dock2*, *Itk*, *Ptprc*) and fibroblasts (*Lab1*, *Col1a1*, *Col1a5*, *Vim*, *Pdgfra*).

**S2 Fig**. **Single-cell RNA sequencing cell populations by Zeitgeber Time**. **A)** Cell population Uniform manifold approximation and projection (UMAP) plot visualization including all cells from each of the time points combined into a single projection. UMAP plot visualization depicts the cells separated by Zeitgeber Time (ZT). This demonstrates that all cell populations are well represented across each time point without preferential isolation in one specific time point compared to the others. B) Shown is the number of cells detected in each cell population at each ZT and the total number of cells for the population across all ZTs.

**S3 Fig**. **nCV analysis pancreas versus liver**. Shown is the normalized coefficient of variation (nCV) for the combined cell populations from the liver single-cell RNA sequencing dataset (i.e., inclusive of all cells) and from normal murine pancreas for each core clock gene (left). The normal murine pancreas was derived from whole pancreas isolated from 4 mice every 4 hours for 48 hours, RNA isolated, and bulk RNA sequencing performed.^19^ The core clock genes demonstrated clear oscillation via meta2d analysis from the pancreatic tissue. Thus, the nCV from the pancreas is indicative of a healthy and robust circadian clock. The liver, which is a similarly metabolic organ, demonstrates no significant difference in nCV compared to the pancreas (p = 0.088, t-test).

**S4 Fig**. **nCV analysis of liver cell populations**. Shown is the normalized coefficient of variation (nCV) for each of the core clock genes in individual cell populations including hepatocytes 1, hepatocytes 2, pericentral hepatocytes, endothelial cells 2, cholangiocytes and lymphocytes.

**S5 Fig**. **Clock correlation analysis of liver cell populations**. Shown are the clock correlation heat maps for various cell populations (the remainder are shown in **Figure 2**) which depict the phase relationship of the core clock genes. Dark red indicates highly positively correlated while dark blue indicates strong negative correlation. The positive arm members (e.g., BMAL1, CLOCK) possess similar Zeitgeber Time of peak phase (Spearman’s rho (ρ) ∼1) whereas positive arm members and negative arm members typically demonstrate antiphasic expression (ρ ∼ -1). Using the set of core clock genes, the correlation of each gene against the others is determined and mapped in matrix form. Z-statistic (z-stat) values represent the extent that these established correlations are preserved in each of the cell populations evaluated, with higher values indicating phase relationships that are closer to the circadian gene atlas baseline (i.e., highly preserved clock correlation).

**S6 Fig**. **Pseudotime trajectory of individual core clock genes**. Shown is a pseudotime trajectory visualizing the expression of the representative individual genes from the E-Box activators (*Npas2*), strong E-Box targets (*Nr1d1*, *Dbp*, *Per1*, *Tef*), and E-Box with other inputs (*Per2*, Cry2, Nfil3, *Bhlhe41*). Zero pseudotime is set to the point of the lowest gene expression. *Bmal1*, *Clock*, *Nr1d2*, *Per3*, *Cry1*, and *Rorc* are shown in **Fig 3**.

**S7 Fig**. **Ordered gene list phase of expression in cell populations**. The top 100 clock driven genes were identified from an existing bulk liver RNA sequencing dataset.^21^ These genes were ordered according to phase of expression via meta2d and gene expression evaluated over time (at each ZT). This was visualized via heatmap for each of the different cell populations. Shown is the heatmap for hepatocytes 1, hepatocytes 2, endothelial cells 2, cholangiocytes, fibroblasts and lymphocytes. The pericentral hepatocytes, periportal hepatocytes, Kupffer cells and endothelial cells 1 are shown in **Fig 5**.

**S8 Fig**. **Phase diagram core clock genes**. The phase of expression of the core clock genes was identified in liver from bulk RNA sequencing (blue).^24^ Additionally, the phase of expression of the core clock genes in the single-cell dataset (all cells, all clusters) was determined via meta2d analysis (purple). These are overlayed on a phase diagram to compare the phase of expression between the two datasets.

**S9 Fig**. **Pseudoreplicate pseudobulk strategy for detecting oscillating gene expression in single-cell datasets**. In the pseudobulk strategy, expression levels of all cells within a cluster are aggregated for each Zeitgeber Time (ZT) to generate one biological replicate for each cluster at each ZT. This would liken the results to bulk RNA sequencing were only that cell population isolated (i.e., pseudobulk). This single biological replicate strategy was used to calculate nCV, clock correlation and core clock gene expression / phase (**Fig 2-5 and S2-7 Fig**). However, this resulted in few detected oscillating genes including lack of detection of rhythmicity in the core clock genes. Consequently, we employed a pseudoreplicate / pseudobulk approach. This entailed randomly partitioning each population of cells (each cluster) into three subsets containing equal numbers of cells, followed by pseudobulking each pseudoreplicate subset. The end result is three pseudoreplicates treated as biological replicates per cell population at each ZT, which enables a much more robust application of rhythmicity detection algorithms. MetaCycle meta2d was used for detection of rhythmically expressed genes in each cell population. To mitigate against false positives, the pseudoreplicate approach was repeated for ten separate random partitions in each cell population (n = 10 trials). Following each trial, meta2d was applied to detect rhythmically expressed genes. Oscillating genes (FDR *q* < 0.05) detected in ten out of ten iterations was the threshold for rhythmicity. Although the figure representation above appears to show discrete parts of the specific cluster for the partitioning, this was actually a random selection of cells distributed throughout the cell population for the pseudoreplicate / pseudobulk approach.

**S10 Fig**. **Oscillating genes by cell cluster and pseudoreplicate iterations**. The pseudoreplicate / pseudobulk strategy was run for ten separate iterations for each cell population and rhythmically expressed genes identified by meta2d analysis (*q* < 0.05). Oscillating genes that were detected in one or more trials (iteration) are depicted in the ‘At least 1 set’ row. As expected, fewer oscillating genes were detected as the cut off for involved iterations increased. The greatest disparities in fraction of oscillating genes were found in cell populations with the fewest number of cells. For example, 186 oscillating genes were found in at least one of ten iterations in cholangiocytes versus 14 oscillating genes with a threshold of ten out of ten iterations, or 7.5%. Meanwhile, 2888 genes were identified in at least one iteration in periportal hepatocytes versus 2068 genes in all ten iterations, or 71.6%. To minimize false positivity, and accepting a trade-off of false negatives, we identified oscillating genes as those that were rhythmically expressed in all ten iterations (meta2d *q* < 0.05).

**S11 Fig**. **UpSet plot oscillating genes by cell cluster**. Shown is an Upset plot with the different individual liver cell populations. The Set Size pertains to the number of oscillating genes identified per cell cluster (meta2d FDR *q* < 0.05 and 10/10 iterations) whereas the Interaction Size depicts the number of genes that are expressed by two or more clusters of cells (dots and line), or the number of genes that are solely expressed by the individual cell cluster (dot).

**S12 Fig**. **Overlapping oscillating gene expression with clock-driven genes**. Shown are Venn diagrams containing the number of oscillating genes detected using the pseudoreplicate / pseudobulk strategy, with a threshold of FDR *q* < 0.05 and rhythmically expressed genes from all ten independent iterations (10/10 pseudoreplicates). Depicted is the overlap in rhythmically expressed genes between known clock-driven genes from a bulk RNA sequencing dataset (bulk RNAseq)^21^ and four representative cell clusters including hepatocyte 1, hepatocyte 2, endothelial cells 1 and pericentral hepatocytes.

**S13 Fig**. **KEGG and Reactome pathway analysis of oscillating genes in endothelial cells**. Clusters with sufficient number of rhythmically expressed genes underwent KEGG and Reactome pathway analysis. Depicted are the top ten pathways from KEGG pathway analysis using the oscillating genes from the endothelial cells 1 cluster. The circle size represents number of genes within the particular pathway and coloration represents adjusted p-value. Heatmaps were also generated depicting the specific oscillating genes identified within selected pathways (representative heatmaps from both Reactome and KEGG). These demonstrate discrete peak phases and trough phases of expression as would be expected for rhythmically expressed genes. Most of the pathways identified from Reactome and KEGG for each cell cluster were shared by more than one cluster, including metabolic pathways (e.g., fatty acid metabolism).

**S14 Fig**. **KEGG and Reactome pathway analysis of oscillating genes in hepatocytes 1 and 2**. Depicted are the top ten pathways from KEGG pathway analysis using the oscillating genes from the hepatocytes 1 cluster (top) and hepatocytes 2 cluster (bottom). The circle size represents number of genes within the particular pathway and coloration represents adjusted p-value. Heatmaps were also generated depicting the specific oscillating genes identified within selected pathways (representative heatmaps from both Reactome and KEGG). These demonstrate discrete peak phases and trough phases of expression, with most of the pathways identified from Reactome and KEGG for each cell cluster shared by several clusters, including metabolic pathways (e.g., PPAR signaling, fatty acid metabolism) and bile acid synthesis / metabolism.

**S15 Fig**. **KEGG pathway and Reactome pathway analysis of oscillating genes for pericentral and periportal hepatocytes**. KEGG pathway and Reactome pathway analyses were performed using the oscillating genes from pericentral hepatocytes (top) and periportal hepatocytes (bottom). Shown are the top ten pathways identified via KEGG analysis where the circle size represents number of genes within the pathway and coloration represents adjusted p-value. Heatmaps were also generated depicting the specific oscillating genes identified within selected pathways (representative heatmaps from both Reactome and KEGG). These demonstrate discrete peak phases and trough phases of expression as would be expected for rhythmically expressed genes. Similar to the other liver cell populations, most of the pathways identified were comprised of metabolic pathways (e.g., PPAR signaling and bile acid synthesis / metabolism). Shown for the periportal hepatocytes cluster is the side by side of KEGG pathway analysis (left) and Reactome pathway analysis (right), for the top ten pathways in each.

**S1 File**. **Average pseudobulk rhythmicity analysis for individual cell populations**.

**S2 File**. **Single-cell replicates rhythmicity analysis for individual cell populations**.

**S3 File**. **Pseudoreplicate trial summary and meta2d analysis**.

**S4 File**. **Trial iteration with significant genes for individual cell populations**.

**S5 File**. **Pathway analysis for individual cell populations**.

**GEO ID for data availability**: GSE292219

## Notes

### Competing Interest Statement

The authors have declared no competing interest.

## References

1. Takahashi JS. Transcriptional architecture of the mammalian circadian clock. Nat Rev Genet. 2017;18(3):164–79.

2. Bass J, Takahashi JS. Circadian integration of metabolism and energetics. Science. 2010;330(6009):1349–54.

3. Panda S. Circadian physiology of metabolism. Science. 2016;354(6315):1008–15.

4. Crosby P, Hamnett R, Putker M, Hoyle NP, Reed M, Karam CJ, et al. Insulin/IGF-1 Drives PERIOD Synthesis to Entrain Circadian Rhythms with Feeding Time. Cell. 2019;177(4):896–909 e20.

5. Fougeray T, Polizzi A, Regnier M, Fougerat A, Ellero-Simatos S, Lippi Y, et al. The hepatocyte insulin receptor is required to program the liver clock and rhythmic gene expression. Cell Rep. 2022;39(2):110674.

6. Guan D, Lazar MA. Interconnections between circadian clocks and metabolism. J Clin Invest. 2021;131(15).

7. Guan D, Xiong Y, Borck PC, Jang C, Doulias PT, Papazyan R, et al. Diet-Induced Circadian Enhancer Remodeling Synchronizes Opposing Hepatic Lipid Metabolic Processes. Cell. 2018;174(4):831–42 e12.

8. Guan D, Xiong Y, Trinh TM, Xiao Y, Hu W, Jiang C, et al. The hepatocyte clock and feeding control chronophysiology of multiple liver cell types. Science. 2020;369(6509):1388–94.

9. Kettner NM, Voicu H, Finegold MJ, Coarfa C, Sreekumar A, Putluri N, et al. Circadian Homeostasis of Liver Metabolism Suppresses Hepatocarcinogenesis. Cancer Cell. 2016;30(6):909–24.

10. Greenwell BJ, Trott AJ, Beytebiere JR, Pao S, Bosley A, Beach E, et al. Rhythmic Food Intake Drives Rhythmic Gene Expression More Potently than the Hepatic Circadian Clock in Mice. Cell Rep. 2019;27(3):649–57 e5.

11. Koronowski KB, Kinouchi K, Welz PS, Smith JG, Zinna VM, Shi J, et al. Defining the Independence of the Liver Circadian Clock. Cell. 2019;177(6):1448–62 e14.

12. Xiong X, Kuang H, Ansari S, Liu T, Gong J, Wang S, et al. Landscape of Intercellular Crosstalk in Healthy and NASH Liver Revealed by Single-Cell Secretome Gene Analysis. Mol Cell. 2019;75(3):644–60 e5.

13. Wen Y, Lambrecht J, Ju C, Tacke F. Hepatic macrophages in liver homeostasis and diseases-diversity, plasticity and therapeutic opportunities. Cell Mol Immunol. 2021;18(1):45–56.

14. Halpern KB, Shenhav R, Matcovitch-Natan O, Toth B, Lemze D, Golan M, et al. Single-cell spatial reconstruction reveals global division of labour in the mammalian liver. Nature. 2017;542(7641):352–6.

15. Xu M, Xu HH, Lin Y, Sun X, Wang LJ, Fang ZP, et al. LECT2, a Ligand for Tie1, Plays a Crucial Role in Liver Fibrogenesis. Cell. 2019;178(6):1478–92 e20.

16. Wen S, Ma D, Zhao M, Xie L, Wu Q, Gou L, et al. Spatiotemporal single-cell analysis of gene expression in the mouse suprachiasmatic nucleus. Nat Neurosci. 2020;23(3):456–67.

17. Xu B, Ma D, Abruzzi K, Braun R. Detecting Rhythmic Gene Expression in Single-cell Transcriptomics. J Biol Rhythms. 2024:7487304241273182.

18. Duan J, Ngo MN, Karri SS, Tsoi LC, Gudjonsson JE, Shahbaba B, et al. tauFisher predicts circadian time from a single sample of bulk and single-cell pseudobulk transcriptomic data. Nat Commun. 2024;15(1):3840.

19. Reinke H, Asher G. Circadian Clock Control of Liver Metabolic Functions. Gastroenterology. 2016;150(3):574–80.

20. Wang S. Protocol to obtain high-quality single-cell RNA-sequencing data from mouse liver cells using centrifugation. STAR Protoc. 2022;3(4):101824.

21. Schwartz PB, Nukaya M, Berres ME, Rubinstein CD, Wu G, Hogenesch JB, et al. The circadian clock is disrupted in pancreatic cancer. PLoS Genet. 2023;19(6):e1010770.

22. Wu G, Francey LJ, Ruben MD, Hogenesch JB. Normalized coefficient of variation (nCV): a method to evaluate circadian clock robustness in population scale data. Bioinformatics. 2021;37(23):4581–3.

23. Wu G, Ruben MD, Schmidt RE, Francey LJ, Smith DF, Anafi RC, et al. Population-level rhythms in human skin with implications for circadian medicine. Proc Natl Acad Sci U S A. 2018;115(48):12313–8.

24. Shilts J, Chen G, Hughey JJ. Evidence for widespread dysregulation of circadian clock progression in human cancer. PeerJ. 2018;6:e4327.

25. Schwartz PB, Walcheck MT, Berres M, Nukaya M, Wu G, Carrillo ND, et al. Chronic jetlag-induced alterations in pancreatic diurnal gene expression. Physiol Genomics. 2021;53(8):319–35.

26. Squair JW, Gautier M, Kathe C, Anderson MA, James ND, Hutson TH, et al. Confronting false discoveries in single-cell differential expression. Nat Commun. 2021;12(1):5692.

27. Abe YO, Yoshitane H, Kim DW, Kawakami S, Koebis M, Nakao K, et al. Rhythmic transcription of Bmal1 stabilizes the circadian timekeeping system in mammals. Nat Commun. 2022;13(1):4652.

28. Finak G, McDavid A, Yajima M, Deng J, Gersuk V, Shalek AK, et al. MAST: a flexible statistical framework for assessing transcriptional changes and characterizing heterogeneity in single-cell RNA sequencing data. Genome Biol. 2015;16:278.

29. Wu G, Anafi RC, Hughes ME, Kornacker K, Hogenesch JB. MetaCycle: an integrated R package to evaluate periodicity in large scale data. Bioinformatics. 2016;32(21):3351–3.

30. Weger BD, Gobet C, David FPA, Atger F, Martin E, Phillips NE, et al. Systematic analysis of differential rhythmic liver gene expression mediated by the circadian clock and feeding rhythms. Proc Natl Acad Sci U S A. 2021;118(3).

31. Heumos L, Schaar AC, Lance C, Litinetskaya A, Drost F, Zappia L, et al. Best practices for single-cell analysis across modalities. Nat Rev Genet. 2023;24(8):550–72.

32. Zadravec D, Brolinson A, Fisher RM, Carneheim C, Csikasz RI, Bertrand-Michel J, et al. Ablation of the very-long-chain fatty acid elongase ELOVL3 in mice leads to constrained lipid storage and resistance to diet-induced obesity. FASEB J. 2010;24(11):4366–77.

33. Michael MD, Kulkarni RN, Postic C, Previs SF, Shulman GI, Magnuson MA, et al. Loss of insulin signaling in hepatocytes leads to severe insulin resistance and progressive hepatic dysfunction. Mol Cell. 2000;6(1):87–97.

34. de Aguiar Vallim TQ, Tarling EJ, Edwards PA. Pleiotropic roles of bile acids in metabolism. Cell Metab. 2013;17(5):657–69.

35. Marinov GK, Williams BA, McCue K, Schroth GP, Gertz J, Myers RM, et al. From single-cell to cell-pool transcriptomes: stochasticity in gene expression and RNA splicing. Genome Res. 2014;24(3):496–510.

36. Chen H, Gao L, Yang D, Xiao Y, Zhang M, Li C, et al. Coordination between the circadian clock and androgen signaling is required to sustain rhythmic expression of Elovl3 in mouse liver. J Biol Chem. 2019;294(17):7046–56.

37. Wang Q, Tikhonenko M, Bozack SN, Lydic TA, Yan L, Panchy NL, et al. Changes in the daily rhythm of lipid metabolism in the diabetic retina. PLoS One. 2014;9(4):e95028.

38. Chen D, Chao DL, Rocha L, Kolar M, Nguyen Huu VA, Krawczyk M, et al. The lipid elongation enzyme ELOVL2 is a molecular regulator of aging in the retina. Aging Cell. 2020;19(2):e13100.

39. Reddy AB, O’Neill JS. Healthy clocks, healthy body, healthy mind. Trends Cell Biol. 2010;20(1):36–44.

40. Schwarz M, Lund EG, Setchell KD, Kayden HJ, Zerwekh JE, Bjorkhem I, et al. Disruption of cholesterol 7alpha-hydroxylase gene in mice. II. Bile acid deficiency is overcome by induction of oxysterol 7alpha-hydroxylase. J Biol Chem. 1996;271(30):18024–31.

41. Aziz H, Underwood PW, Gosse MD, Afyouni S, Kamel I, Pawlik TM. Hepatic adenoma: evolution of a more individualized treatment approach. J Gastrointest Surg. 2024;28(6):975–82.

42. Hao Y, Hao S, Andersen-Nissen E, Mauck WM, 3rd, Zheng S, Butler A, et al. Integrated analysis of multimodal single-cell data. Cell. 2021;184(13):3573–87 e29.

43. MacParland SA, Liu JC, Ma XZ, Innes BT, Bartczak AM, Gage BK, et al. Single cell RNA sequencing of human liver reveals distinct intrahepatic macrophage populations. Nat Commun. 2018;9(1):4383.

44. Germain PL, Lun A, Garcia Meixide C, Macnair W, Robinson MD. Doublet identification in single-cell sequencing data using scDblFinder. F1000Res. 2021;10:979.

45. Choudhary S, Satija R. Comparison and evaluation of statistical error models for scRNA-seq. Genome Biol. 2022;23(1):27.

46. McInnes LH, John; Melville, James. UMAP: Uniform Manifold Approximation and Projection for Dimension Reduction. 2018.

47. Conway JR, Lex A, Gehlenborg N. UpSetR: an R package for the visualization of intersecting sets and their properties. Bioinformatics. 2017;33(18):2938–40.

48. Larsson JG, A Jonathan R.; Gustafsson, Peter; Eberly, David H.; Huber, Emanuel; Prive, Florian. Area-proportional euler and venn diagrams with ellipses. Version 7.0.2 ed: CRAN; 2024.

49. Xu S, Hu E, Cai Y, Xie Z, Luo X, Zhan L, et al. Using clusterProfiler to characterize multiomics data. Nat Protoc. 2024;19(11):3292–320.

50. Gene Ontology C, Aleksander SA, Balhoff J, Carbon S, Cherry JM, Drabkin HJ, et al. The Gene Ontology knowledgebase in 2023. Genetics. 2023;224(1).

51. Ashburner M, Ball CA, Blake JA, Botstein D, Butler H, Cherry JM, et al. Gene ontology: tool for the unification of biology. The Gene Ontology Consortium. Nat Genet. 2000;25(1):25–9.

52. Kanehisa M, Goto S. KEGG: kyoto encyclopedia of genes and genomes. Nucleic Acids Res. 2000;28(1):27–30.

53. Milacic M, Beavers D, Conley P, Gong C, Gillespie M, Griss J, et al. The Reactome Pathway Knowledgebase 2024. Nucleic Acids Res. 2024;52(D1):D672–D8.

